# Understanding the relationship between dispersal and range size

**DOI:** 10.1101/2021.09.29.462346

**Authors:** Adriana Alzate, Renske E. Onstein

**Affiliations:** German Centre for Integrative Biodiversity Research (iDiv) Halle-Jena-Leipzig, Puschstr. 4, D-04103 Leipzig, Germany; Leipzig University, Ritterstraße 26, 04109 Leipzig, Germany

## Abstract

Understanding what drives the vast variability in species range size is still an outstanding question. Among the several processes potentially affecting species ranges, dispersal is one of the most prominent hypothesized predictors. However, the theoretical expectation of a positive dispersal-range size relationship has received mixed empirical support. Here, we synthesized results from 84 studies to investigate in which context dispersal is most important in driving species range size. We found that dispersal traits – proxies for dispersal ability – explain range sizes more often in temperate and subtropical regions than in tropical regions, when considering multiple components of dispersal, and when investigating a large number of species to capture dispersal and range size variation. In plants, positive effects of dispersal on range size were less often detected when examining broad taxonomic levels. In animals, dispersal is more important for range size increase in ectotherms than in endotherms. Our synthesis emphasizes the importance of considering different aspects of the dispersal process -departure, transfer, settlement-, niche aspects and evolutionary components, like time for range expansion and past geological-environmental dynamics. We therefore call for a more integrative view of the dispersal process and its causal relationship with range size.

## Introduction

Species geographical range is a fundamental unit in macroecology and is a main predictor of extinction risk across organisms (Brown et al. 1996, Purvis et al. 2000, Chichorro et al. 2019). As the distribution of a species provides important information on their ecology and evolution, understanding what drives the vast variability in species range size has for long been of interest to paleontologist, biogeographers, macroecologists and evolutionary biologists (Brown et al. 1996, Gaston 2003, Gaston 2009, Gaston & Fuller 2009). Several mechanisms might underlie species geographical ranges, such as environmental and physical constraints, differences in niche requirements, population abundance, latitudinal gradients, differences in body size or trophic level, colonization-extinction dynamics, species age and dispersal ability (Gaston 2003). Although all these mechanisms can simultaneously interact to produce the empirical range sizes, dispersal has received the most interest, likely because it is one of the most prominent processes affecting range size (Hanski et al. 1993, Lester et al. 2007, Sheth et al. 2020). However, despite major efforts to link dispersal ability to range size, the theoretical expectation of a positive dispersal-range size relationship has received mixed empirical support both among and within taxa (Lester et al. 2007, Luiz et al. 2013, Alzate et al. 2019a, Sheth et al. 2020).

Here we define dispersal as any movement of individuals or propagules potentially leading to gene flow across space (Ronce 2007, Clobert et al. 2012). Both theoretical and experimental work have shown that dispersal can positively affect the mechanisms that allow attaining large geographical ranges (Holt 2003, Holt et al. 2005, Sexton et al. 2009, Alzate et al. 2019b). For instance, dispersal promotes range expansion by facilitating the colonization of new habitats and promoting local adaptation (Holt & Gomulkiewicz 1997, Alzate et al. 2019b). Dispersal also prevents range contraction by decreasing extinction risk and allowing populations to persist in suboptimal habitats (Hufbauer et al. 2015, Alzate et al. 2019b), as it provides demographic (Brown & Kondric-Brown 1977) and genetic (Holt & Gomulkiewicz 1997) rescue. Furthermore, simulation models (spatial explicit neutral models) predict a positive effect of dispersal on species range size (Rangel-Diniz-Filho 2005, Alzate et al. 2019c). An outstanding question is therefore why positive effects of dispersal on range size are found in some cases, but not in others.

Discrepancies between expected and observed dispersal-range size relationships might emerge for different reasons. Firstly, studies differ in their dispersal ‘definitions’ and therefore also differ in which phenotypic traits are considered to be associated to dispersal ability. Which dispersal-related traits to choose is not a trivial question, particularly because dispersal and dispersal distance are emergent properties (complex traits) resulting from the interactive effects of various dispersal-related traits (e.g., morphology, physiology, behavioural and life history traits), which are highly dependent on the type of organism studied (Ronce 2007, Bonte et al. 2012, Matthysen 2012, Ronce & Clobert 2012, Travis et al. 2012, Sheth 2020, Green et al. 2021). Moreover, dispersal can occur at different life stages (e.g., seeds, eggs, juveniles, adults), it is composed by three phases including departure (i.e., decision to leave the old habitat), transfer (i.e., displacement from the old to a new habitat) and settlement (i.e., arrival and settlement in the new habitat), and can be shaped by external factors and evolve (Ronce 2007, Matthysen 2012). Although dispersal kernels are in general good proxies for dispersal abilities across species, there are still difficulties to measure the tail of the kernel, which is of major importance when scaling up to distribution dynamics (Clobert et al. 2012). Therefore, different measures of dispersal may capture different components of the dispersal process, and may thus affect the resulting dispersal-range size relationship.

Secondly, true biological differences between study organisms might determine how dispersal correlates with range size. For instance, it is possible that the dispersal-range size relationship differs between active (e.g., reptiles, birds, mammals) and passive dispersers (e.g., plants, diatoms, marine molluscs), because dispersal (the transfer phase) for passive dispersers is outside the control of the individual, as it depends on external forces (e.g., currents, wind, gravity, other organisms) with a high stochastic component (Matthysen 2012). Moreover, the nature of the dispersal medium of marine and terrestrial realms can lead to differences in dispersal ability between terrestrial and marine organism. In marine systems, passive rather than active dispersal is favoured, which may lead to fewer dispersal-related adaptations in marine than in terrestrial systems (Burgess et al. 2005). The higher connectivity between marine habitats may also erase the link between dispersal ability and range sizes (Mora et al. 2012). In terrestrial systems, dispersal may be more difficult without specialised adaptations (such as wings to fly, fleshy fruits to attract animal dispersers, etc.) (Burgess et al. 2015). In addition, human activities can uncouple the dispersal-range size relationship by decreasing species distributions (Webb & Gaston 2000). This might be more common on terrestrial than in marine systems, where large scale human impact has a longer history. Similarly, dispersal may be less limiting in endotherms than ectotherms, because endotherms possess broader thermal niches and higher thermal tolerances due to high metabolic rates compared to ectotherms, facilitating settlement. Even though endo- and ectotherms might not differ in the dispersal capacities required during the transfer phase of dispersal, endothermy might have an advantage during the settlement phase. Thus, physiological tolerance rather than dispersal ability may in some cases be the limiting factor when it comes to range size (Pie et al. 2021).

Thirdly, evolutionary history may affect the current dispersal-range size relationship by determining time and potential for (past) range expansion, or the evolution of dispersal-related traits that facilitated past long-distance dispersal (Onstein et al. 2019). For example, range size is likely to vary with species age because species need time to expand their range (Willis 1922, Webb & Gaston 2000, Gaston 2003), and the dispersal-range size relationship may thus be obscured when studying species of different ages. Furthermore, past climate changes (e.g., since the last glacial maximum), population connectivity, and availability of suitable settlement environments, may have affected the rate of range size expansion (e.g., ‘Reid’s Paradox’), and may explain distinct dispersal-range size relationships across biogeographical realms and climate zones that differ in their glaciation history, for example (Svenning & Skov 2004, Svenning et al. 2008).

Lastly, besides biological and evolutionary reasons, intrinsic differences between studies, such as study design, methodology, data, or analytical approach might explain absence of a relationship between dispersal and range size. Studies examining how dispersal affects range size differ in taxonomic scope (e.g., ‘genus’, ‘family’, ‘phylum’), taxonomic unit of analysis (e.g., at ‘genus’ or ‘species’ level), the measure of species range size (e.g., ‘extent of occurrence’ or ‘area of occupancy’), range completeness (e.g., ‘partial’ or ‘complete’ measures of range size), and the number of considered species and dispersal-related traits to capture variation in the dispersal-range size relationship. For example, range size measures can under- or overestimate true range sizes by including or excluding discontinuities in the spatial distribution of taxa. Similarly, high taxonomic units of analysis (e.g., ‘genus’ or ‘family’ instead of ‘species’) ignores within-clade variation in dispersal. The benefit of using comprehensive data, that is, bigger areas that include complete ranges, over partial data (i.e., smaller areas that do not include complete ranges) has been shown for understanding the body size - range size relationship in animals, with more consistently positive relationships when using comprehensive data (Gaston & Blackburn 1996).

Despite all possible caveats and warnings about several of these problems (Blackburn & Gaston 1998, Alzate et al. 2019a, Johnson et al. 2021), a comprehensive methodological framework to study dispersal and range size is missing. Here we performed a systematic review to investigate the causes of variation in the dispersal-range size relationship by collating 478 dispersal-range size relationships from 86 independent studies. Firstly, we investigated and synthesized the spread of evidence for the dispersal-range size relationship between regions, realms and clades. Secondly, we quantified how differences between studies regarding dispersal and range size characteristics related to transfer, settlement and evolution, and potential methodological differences (range size definitions, spatial corrections and taxonomy) can affect the overall dispersal-range size relationship (Table 1). Finally, we discuss these results in the context of the complexity of the dispersal process – from departure, to transfer, to settlement.

**Table 1.**
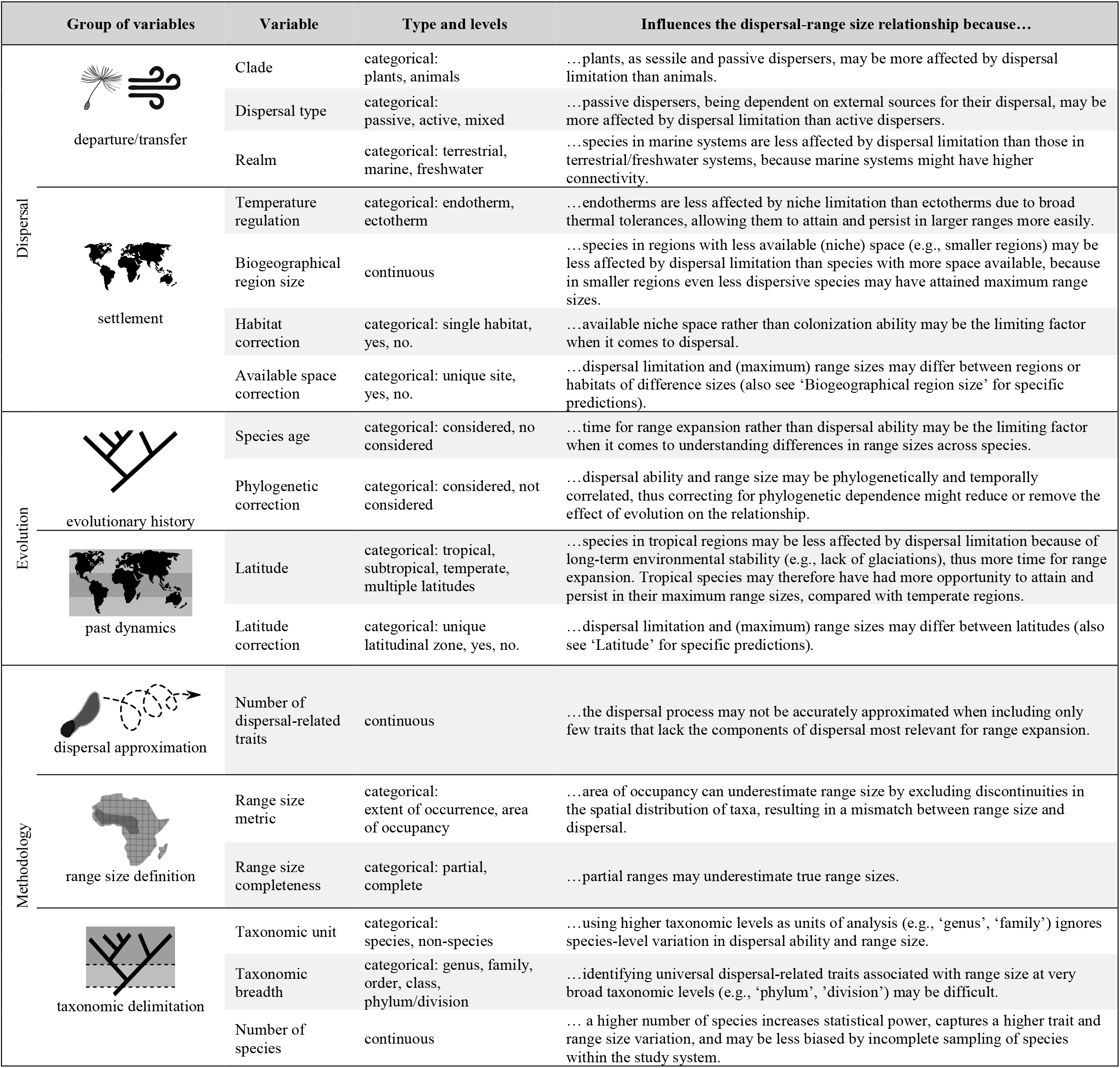
Moderator candidates of the dispersal-range size relationship as synthesized in this study. Variables were classified into six groups depending on their hypothesized effect on the dispersal-range size relationship: departure/transfer variables that directly affect movement/transfer of species, settlement variables or corrections that influence the potential and realized niche space, time variables or corrections related to evolutionary history and past dynamics that may influence range size, and three methodological type variables that may bias the inference of the dispersal-range size relationship. Description of each variable and the prediction of why and how it potentially influences the dispersal-range size relationship is provided.

|  | Group of variables | Variable | Type and levels | Influences the dispersal-range size relationship because... |
| --- | --- | --- | --- | --- |
| Dispersal | departure/transfer | Clade | categorical: plants, animals | ...plants, as sessile and passive dispersers, may be more affected by dispersal limitation than animals. |
|  |  | Dispersal type | categorical: passive, active, mixed | ...passive dispersers, being dependent on external sources for their dispersal, may be more affected by dispersal limitation than active dispersers. |
|  |  | Realm | categorical: terrestrial, marine, freshwater | ...species in marine systems are less affected by dispersal limitation than those in terrestrial/freshwater systems, because marine systems might have higher connectivity. |
|  | settlement | Temperature regulation | categorical: endotherm, ectotherm | ...endotherms are less affected by niche limitation than ectotherms due to broad thermal tolerances, allowing them to attain and persist in larger ranges more easily. |
|  |  | Biogeographical region size | continuous | ...species in regions with less available (niche) space (e.g., smaller regions) may be less affected by dispersal limitation than species with more space available, because in smaller regions even less dispersive species may have attained maximum range sizes. |
|  |  | Habitat correction | categorical: single habitat, yes, no. | ...available niche space rather than colonization ability may be the limiting factor when it comes to dispersal. |
|  |  | Available space correction | categorical: unique site, yes, no. | ...dispersal limitation and (maximum) range sizes may differ between regions or habitats of difference sizes (also see 'Biogeographical region size' for specific predictions). |
| Evolution | evolutionary history | Species age | categorical: considered, no considered | ...time for range expansion rather than dispersal ability may be the limiting factor when it comes to understanding differences in range sizes across species. |
|  |  | Phylogenetic correction | categorical: considered, not considered | ...dispersal ability and range size may be phylogenetically and temporally correlated, thus correcting for phylogenetic dependence might reduce or remove the effect of evolution on the relationship. |
|  | past dynamics | Latitude | categorical: tropical, subtropical, temperate, multiple latitudes | ...species in tropical regions may be less affected by dispersal limitation because of long-term environmental stability (e.g., lack of glaciations), thus more time for range expansion. Tropical species may therefore have had more opportunity to attain and persist in their maximum range sizes, compared with temperate regions. |
|  |  | Latitude correction | categorical: unique latitudinal zone, yes, no. | ...dispersal limitation and (maximum) range sizes may differ between latitudes (also see 'Latitude' for specific predictions). |
| Methodology | dispersal approximation | Number of dispersal-related traits | continuous | ...the dispersal process may not be accurately approximated when including only few traits that lack the components of dispersal most relevant for range expansion. |
|  | range size definition | Range size metric | categorical: extent of occurrence, area of occupancy | ...area of occupancy can underestimate range size by excluding discontinuities in the spatial distribution of taxa, resulting in a mismatch between range size and dispersal. |
|  |  | Range size completeness | categorical: partial, complete | ...partial ranges may underestimate true range sizes. |
|  | taxonomic delimitation | Taxonomic unit | categorical: species, non-species | ...using higher taxonomic levels as units of analysis (e.g., 'genus', 'family') ignores species-level variation in dispersal ability and range size. |
|  |  | Taxonomic breadth | categorical: genus, family, order, class, phylum/division | ...identifying universal dispersal-related traits associated with range size at very broad taxonomic levels (e.g., 'phylum', 'division') may be difficult. |
|  |  | Number of species | continuous | ... a higher number of species increases statistical power, captures a higher trait and range size variation, and may be less biased by incomplete sampling of species within the study system. |

### Literature search methodology

We conducted a systematic search for studies that examined the relationship between dispersal-related traits and range size. We used Web of Science and the Core Collection database with the key words search criteria “dispers*” OR “dispers* trait” AND “range size” OR “species range” OR “geogra* range”. We restricted our search to scientific articles published in English. Our search yielded a total of 3,139 scientific articles. After a first screening based on article’ titles and abstracts, we discarded studies that were clearly irrelevant (e.g., studies in other fields like physics, studies that do not formally assess the dispersal - range size relationship, studies on single species, studies that use genetic metrics as proxies of dispersal, studies on range expansion or range shifts). We assessed 104 articles for eligibility by reading the full-text and we further excluded perspectives and review articles or studies that did not include empirical tests (i.e., strictly theoretical papers; see Table S1-S4 in Database). After cross-referencing and including newly published articles, 86 studies and 478 relationships between dispersal-related traits and range size were included in our systematic review. For an overview of the search methodology, see the Prisma diagram (Fig. S1 Suppl. Mat.).

### Data extraction and collection

For each relationship reported in the studies, we gathered information on 17 moderators potentially responsible for differences in the dispersal - range size relationship across studies (Table 1). We classified these moderators into six groups of variables related to: 1) departure/transfer stage of dispersal, 2) settlement stage of dispersal, 3) evolution, 4) dispersal approximation, 5) range size definitions, and 6) taxonomic delimitation (see Table 1 for the description of the variables and predictions).

To assess which traits may be most suitable to approach the dispersal process and are most relevant for increasing range size, we reported the effect of each dispersal-related trait included in each model examining the dispersal-range size relationship (Table 2). We classified this trait overview by realm (terrestrial vs. marine) and taxon (clade), due to potential differences between these systems with respect to dispersal-limiting factors (Table 1) and thus their relationship of dispersal-related traits to range size (Table 2). To examine the effect of the moderators on the dispersal-range size relationship for each relationship, we reported the overall effect of dispersal on range size (see Table S5 in Database). When the study reported two or more dispersal-related traits with opposite effects on range size, the overall effect was treated as ‘neutral’. When the study included several dispersal-related traits, some with neutral effects and some with positive effects, we treated the overall effect as ‘positive’, while we treated the overall effect as ‘negative’ when the opposite happened.

**Table 2.**
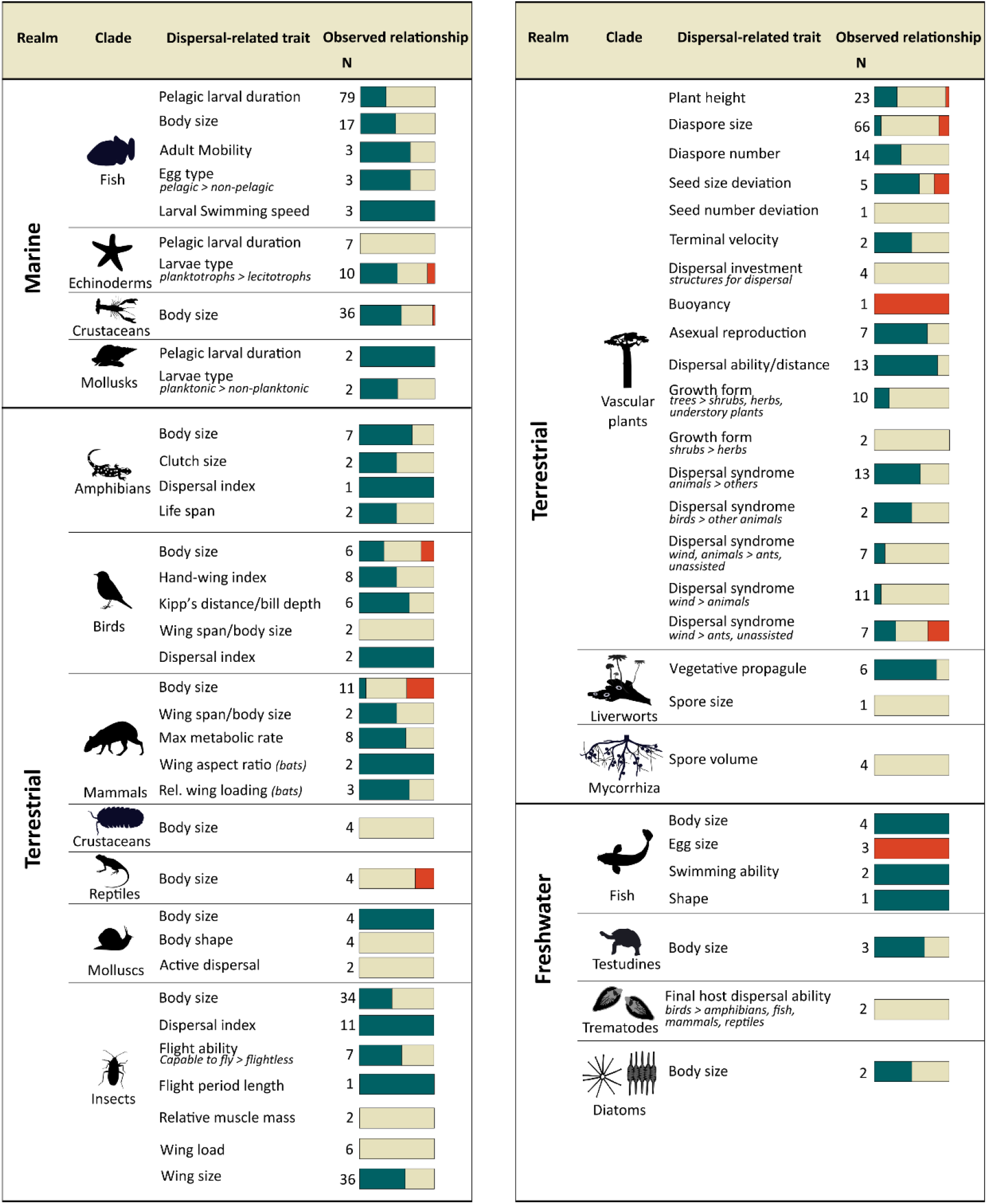
Summary of the observed relationship between dispersal-related traits and range size based on 86 studies and 478 relationships. The relative proportion of the direction of the relationships is indicated (positive: green, neutral: brown, negative: red) for each realm (marine, terrestrial, freshwater) and taxon or clade (fish, echinoderms, crustaceans, molluscs, amphibians, birds, mammals, insects, vascular plants, liverworts, mycorrhiza, testudines, trematodes, diatoms). The number of relationships assessed for each taxon is indicated with *N*. In case of categorical variables, the direction of the trait states is indicated below the trait, e.g., ‘pelagic > non-pelagic’ means that species with pelagic eggs have larger range sizes than those with non-pelagic eggs, and this directional relationship was found in 2 out of 3 cases.

For the calculation of biogeographical region size (which may influence the opportunity for range expansion due to available area) we used GIS layers (Spalding et al. 2007, Olson et al. 2004, and Natural Earth). Among the methodological features, the number of dispersal-related traits refers to the number of traits included as independent factors (e.g., body size, diaspore size, plant height) or combined in a single factor as a complex metric of dispersal (e.g., hand-wing index, PCA axis, relative measurements). For instance, hand-wing index is used as a single factor that is composed of two dispersal-related traits (wing length and first-secondary length). Taxonomic breadth, the lowest taxonomic level that includes all species in the study, was assigned based on the species lists provided in the studies. Relationships using ‘family’ and ‘superfamily’ taxonomic levels were recoded as ‘family’. Similarly, ‘order’ and ‘suborder’ were recoded as ‘order’, and ‘phylum’, ‘division’, ‘subphylum’ and ‘subkingdom’ were recoded as ‘phylum-division’ level. 13 relationships could not be assigned to a particular taxonomic breadth, thus excluded from statistical analyses.

### Moderators affecting the dispersal-range size relationship

To investigate how differences between dispersal processes, evolutionary history, and study design affect the overall dispersal-range size relationship, we fitted a Generalized Linear Mixed Model (GLMM) with Binomial error distribution (link probit) and study ID as a random effect. To test the effect of the 17 moderators (Table 1) on the presence or absence of a dispersal-range size relationship, we performed a forward model selection procedure. We started by fitting a model with only random effects, and sequentially added significant fixed factors until reaching a final model. To decide which variable to include in the model in every time step, we tested for the significance of all fixed factors that can be added to the base model (i.e., all factors that are not already included in the model at that point) using a log-likelihood ratio test to identify the most significant variable to add (and *p* < 0.05). Each model was tested for collinearity using the *vif* function from the ‘car’ R package (Fox & Weisberg 2019). If a new added variable was collinear with one already selected in the model, the new variable was excluded from the selection procedure. We retained variables with a generalized variance inflation factor (GVIF) smaller than 2, which is the square root of the threshold value for the standard VIF (VIF = 5), and indicates limited collinearity. We used generalized linear mixed-effects models in a Bayesian setting (*bglmer* function from R package ‘blme’; Dorie et al. 2021).

We only included positive and neutral relationships as response variables, negative relationships were excluded as they were too few to perform an analytical test on (10 out of 478 relationships in total). All continuous variables (number of dispersal-related traits, biogeographical region size and number of species) were rescaled from 0 to 1. Also, 13 relationships for clades that were only assessed in a single study (mycorrhizal fungal, diatoms and liverworts) were excluded as they could not be placed in any of the two broad clades (animals, vascular plants) assessed here. 36 relationships were also excluded as did not report number of species and 13 were excluded as they missed information on highest taxonomic level. The final model included in total 410 relationships from 81 studies.

In addition to a global model including all relationships, we ran separate analyses for the two broad clades: animals and vascular plants. In animals, we also included the specific taxon (e.g., birds, mammals, fish, insects) as a factor related to the departure/transfer dispersal process, as these clades may differ in their dispersal ability based on certain biological features (e.g., presence or absence of wings). We then excluded invertebrates and trematodes, the former because it cannot be assigned to a specific taxonomic group and the latter, because it only has two reported relationship from a single study (Thieltges et al. 2011). Because temperature regulation and latitude were correlated, we ran two separated models including either temperature regulation or latitude. We recoded ‘tropical’ and ‘multiple latitude’ as ‘tropical-multiple latitude’, and ‘subtropical’ and ‘temperate’ as ‘subtropical-temperate’ for the model including latitude, due to complete separation. In plants, we excluded factors that lacked variation within plant studies, such as dispersal type (passive vs. active), temperature regulation, realm (only studies on terrestrial plants), species age (only one study included this), taxonomic unit, and the taxonomic breadth level ‘class’. Latitude was recoded similarly as done for animals. Model fitting and selection for both subsets was done as described above for the complete data-set.

#### General patterns

Most of the studies assessing the dispersal-range size relationship have focused on marine and terrestrial systems (48 and 31 studies, respectively), whereas freshwater systems have received much less attention (only 7 studies; Fig. 1,2). Regarding taxa, most of studies have focused on vascular plants, fishes, and insects (24, 17 and 14 studies respectively; Fig. 1), whereas dispersal-range size relationships in bryophytes (liverworts), diatoms, trematodes and mycorrhiza fungal (all with a single study) have barely been studied. The majority of studied relationships (55%) showed a neutral effect of dispersal on range size, 40% of relationships were positive and only few (5%) were negative (Fig. 2). While for most of the taxa we did not find a consistent positive association between range size and dispersal, for molluscs, amphibians and birds we found more often positive effects of dispersal on range size than neutral effects (Fig. 1). Marine, terrestrial and freshwater realms showed similar proportions of positive and neutral relationships (Chi-squared test = 0.078, p = 0.67, Fig. 1). Interestingly, plants showed a lower proportion of positive relationships than animals (Chi-squared test = 10.72, p = 0.001, Fig. 1).

**Fig. 1.**
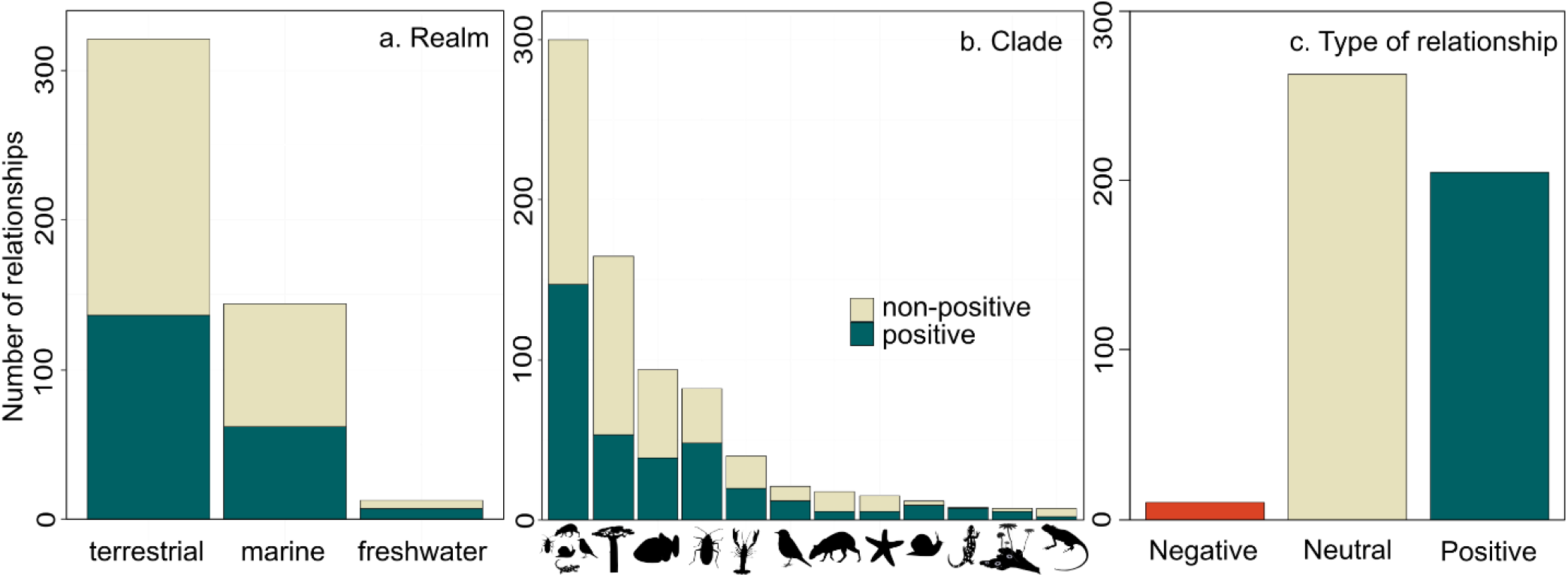
Overview of positive and non-positive relationships between dispersal-related traits and range size examined in this review. The number of positive and non-positive relationships is indicated for (a) each realm (terrestrial, marine, freshwater), (b) clade and specific taxonomic groups (animals, plants, fish, insects, crustaceans, birds, mammals, echinoderms, molluscs, amphibians, liverworts, reptiles) and (c) total. Mycorrhiza, trematodes, diatoms and invertebrates were not included here because they had less than four reported relationships (all non-positive).

**Fig. 2.**
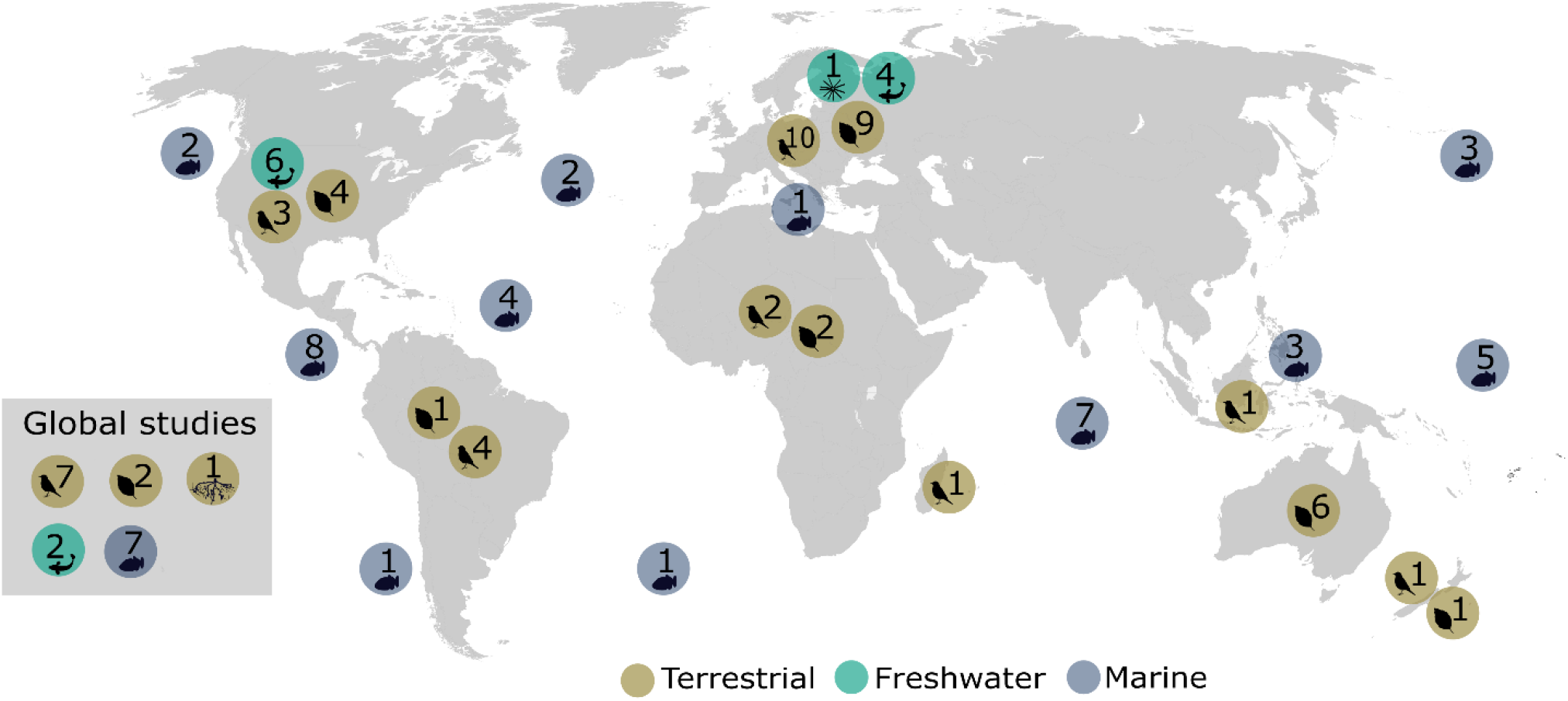
Geographical distribution of studies examining the effect of dispersal on range size. We evaluated this for different clades: animals (fish and bird silhouettes), plants (leaf silhouette), diatoms (diatom silhouette) and mycorrhiza (root silhouette), and realms: terrestrial (brown), freshwater (green) and marine (blue). Except for global studies, studies that were carried out in more than one region are included in each region separately. Geographic location is separated according to the geographical region the study was carried out in, following the Wallace classification for terrestrial and freshwater organisms and marine ecoregions of the world-MEOW for marine organisms.

There is also a clear spatial structure in study location (Fig. 2). For terrestrial systems, most of the studies have been performed in the palearctic region (23 studies), followed by studies including multiple regions (i.e., studies that include more than one region, including global studies) (15 studies), whereas much less attention has been given to neotropical and paleotropical regions (including the Madagascar region). For marine systems, 32% of the studies (10 studies) examined multiple regions, followed by studies in the Indo-Pacific and the Tropical Eastern Pacific (26%, 8 studies each), whereas the Atlantic and Mediterranean regions were much less studied (4 and 1 study respectively, Fig. 1). For freshwater systems, only 2 studies have been performed for multiple regions, the remaining 5 studies have only considered Nearctic a Palearctic regions (2 and 3 studies, respectively).

### Dispersal-related traits

Studies used a wide range of traits as proxies for dispersal ability, which, unsurprisingly, vary substantially with the taxon and system studied (Table 2). For marine animals, traits that allow dispersal during the larval stage (which is often pelagic) are most commonly used. Type of larvae (e.g., pelagic vs. non-pelagic) and egg (e.g., pelagic vs. non-pelagic), and the duration of the pelagic larval stage have generally been considered as proxies of dispersal for fishes, echinoderms and molluscs. Body size has also been used as a proxy for dispersal for crustaceans and fishes. For terrestrial animals, body size has been the main trait used as a proxy for dispersal ability. In addition, traits related to flight ability have been used in taxa for which flight is present (e.g., birds, bats, insects), either as a binary trait (presence vs. absence of wings) or a continuous variable (hand-wing index, wing load or size) reflecting flight potential (Table 2). Life history traits (e.g., clutch size, life span) have mostly been used in amphibians, whereas metabolic rate has been used as a dispersal proxy in mammals. For vascular plants, proxies of dispersal are related to the characteristics of the diaspore (e.g., seed size and number) and dispersal mechanism (i.e., dispersal syndrome, such as wind, water or animal). In freshwater animals, body size is also a common proxy of dispersal (for fish, testudines and diatoms). Dispersal in parasitic trematodes is assumed to be linked to the dispersal ability of the host (e.g. fish, mammals, birds). See Appendix 1 in Suppl. Mat. for an extended description.

### Moderators of the dispersal-range size relationship

The final model, including both plants and animals, explained 41% of the variation (50% considering random effects) and identified four moderators as main factors explaining the differences in the dispersal-range size relationship between studies (Fig. 3). Specifically, studies that included multiple dispersal-related traits, that were carried out in temperate or subtropical regions or that included a larger number of species, showed significantly more positive dispersal-range size relationships than studies including fewer dispersal-related traits, that were carried out in tropical regions (or multiple regions), or that included fewer species in the analysis (Fig. 3, Table S2, S3 Suppl. Mat.). Furthermore, studies carried out at higher taxonomic breadth, such as ‘phylum’ or ‘division’ showed fewer positive relationships between dispersal and range size than studies carried out for a particular ‘class’, ‘order’ or ‘family’.

**Fig. 3.**
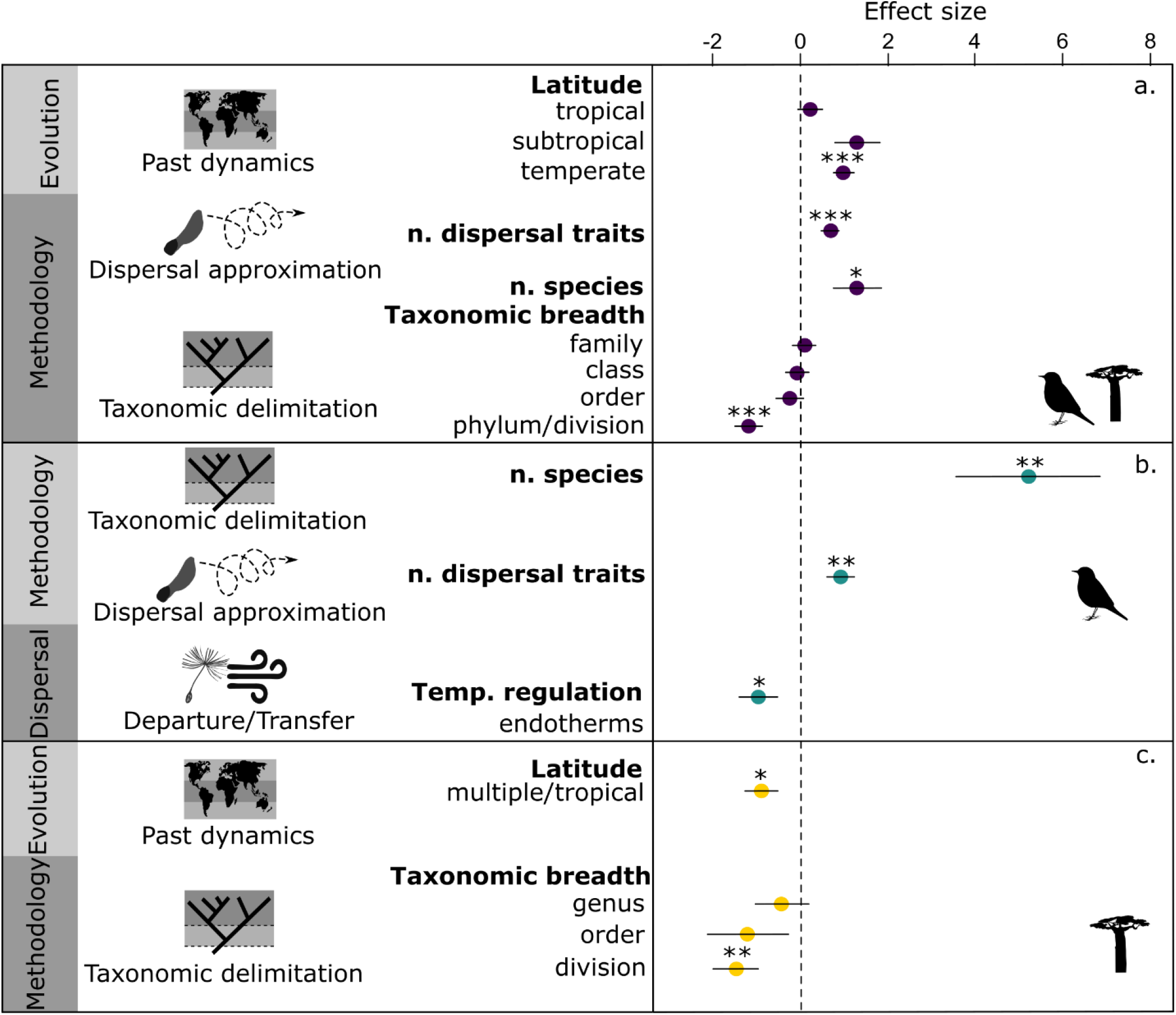
Determinants of the dispersal-range size relationship. Outcomes of the global model (a) and models for animals (b) and plants (c) examining the effect of 19 variables (moderators) on the dispersal-range size relationship across 83 studies and 478 relationships. We show the inferred effect size of standardize predictor variables that remained significant in the final GLMM models (or GLM for plants). Reference levels for the categorical variables are in (a) multiple latitudes (for latitude, i.e. vs. tropical, subtropical, temperature) and class (for taxonomic breadth, i.e. vs. family, genus, order, phylum/division); in (b) area for range size measurement (vs. units/categories/linear/percentage) and ectotherm for temperature regulation (vs. endotherm); and in (c) family for taxonomic breadth (vs. family, class, division). Standard errors around the mean estimates and significance level (* <0.01, ** <0.001, *** <0.0001) of the effect are indicated. n. = ‘number of’.

When examining the relationship between dispersal and range size for animals and plants separately, we found that the factors responsible for a positive dispersal-range size relationship differed between those broad clades (Fig. 3, Table S1, S2 Suppl. Mat.). In animals, similar results as for the global dataset were found, with the number of dispersal-related traits and number of species positively contributing to the dispersal-range size relationship. In addition, we found that studies on endotherms showed fewer significant, positive associations between dispersal and range size than on ectotherms (Fig. 3, Table S1, S2 Suppl. Mat.). In plants, taxonomic breadth remained important as a variable to explain the positive dispersal-range size relationship, with studies carried out at ‘division’ level or higher showing fewer positive dispersal-range size relationships than studies carried out at the ‘family’ level (Fig. 3, Table S1, S2 Suppl. Mat.).

It should be noted that a large sample size may increase the probability of detecting false positives, which could explain our finding that number of species and number of dispersal-related traits affects the dispersal-range size relationship. Although, theoretically, this could go in both a positive or negative direction in terms of dispersal-range size relationships. The distribution of species numbers used in the studies is strongly right-skewed: more than 90% of the studied relationships included less than 500 species (380 relationships, Fig S2 Suppl. Mat.). We performed a sensitivity analysis to explore the effect of outliers for both the number of species and number of dispersal-related traits on our findings. Nevertheless, the positive effect of species number on the dispersal-range size relationship remains when including only studies with less than 500 species, but not when only including studies with less than 100 species (Table S3 Suppl. Mat.). Importantly, the positive effect of including multiple dispersal-related traits remained after excluding possible outliers (studies with 10 dispersal-related traits; Table S3 Suppl. Mat.). Alternatively, our result may reflect serendipity, because the more traits a study includes, the higher the chance becomes that one of those traits relates to range size. We tested whether this is the case by comparing the relationship between the overall dispersal-range size relationship and the number of dispersal-related traits vs. the relationship between the overall dispersal-range size relationship and the number of dispersal ‘factors’ (one factor can be composed of several dispersal-related traits) included in the models (Table S4 Suppl. Mat.). Interestingly, although the effect of the number of dispersal-related traits on the dispersal-range size relationship is significantly positive, the number of dispersal ‘factors’ included in each relationship does not have a significant effect (Table S4 Suppl. Mat.). This means that using multiple dispersal-related traits likely results in a better approximation of the dispersal process and/or a higher probability to capture traits that are relevant for dispersal, increasing the chance to find a positive dispersal-range size relationship.

### The number of species effect on the dispersal - range size relationship

Including a large number of species is advantageous for various reasons. Besides increasing the statistical power of the study, including more species may also capture a larger variation in both dispersal-related traits and range sizes essential to capture the relationship. The number of species used to examine the dispersal-range size relationship across studies ranged from 5 (for reef fishes; Zapata & Herrón 2003 and Lester & Ruttenberg 2005) to 10,338 (for birds; Sheard et al. 2020). A relatively small sample size (i.e., few species investigated), can result from a) study objective, such as the focus on a particular genus that comprises few species, or from b) data limitations, e.g. because trait data are only available for a subset of the species of interest. While data limitations are understandable because it is often difficult to gather information for all species in a large, wide-spread, species-rich clade, it may also come with the risk that the subsample of species and traits are biased and thus not representative for our understanding of the relationship between dispersal and range size. This may lead to biased outcomes (Alzate et al. 2019a), even after data imputation to resolve data gaps (Johnson et al 2021). It is noteworthy that a small sample size might be sufficient if the species pool is also small, so the level of completeness might be the critical issue here. However, we were not able to evaluate this because most studies did not report on sampling completeness.

### The number of dispersal-related traits effect on the dispersal - range size relationship

Our results also indicated that including multiple dispersal-related traits increased the probability of finding a positive dispersal-range size relationship. However, 80% of the studied relationships (383/476) included only a single dispersal trait. As dispersal is an emergent process that results from the combined effects of multiple dispersal characteristics (Ronce 2007, Bonte et al. 2012), including multiple traits may capture the complexity of dispersal better, and thus increase the probability of finding a positive dispersal-range size relationship.

### The biogeographical effect on the dispersal-range size relationship

We also found that studies performed exclusively in temperate or subtropical regions showed more positive associations between dispersal-related traits and range size than studies performed exclusively in the tropics or that included multiple latitudes (Fig. 3). This suggests that (current) species range sizes in tropical organisms may be more independent from dispersal than in subtropical and temperate regions, and supports the hypothesis (Table 1) that species in tropical regions may have had more opportunity to attain and persist in their maximum range sizes compared to temperate species, because range sizes have not been altered as much by past environmental change (e.g., during the Quaternary period, such as lack of glaciations) (Svenning & Skov 2004) and time for range expansion. Indeed, if species in temperate regions are still spreading from their last glacial maximum refugia, their ranges might be smaller than their potential ranges (Svenning et al. 2008), and dispersal-related traits might be more directly affecting range sizes compared to those in tropical regions.

Alternatively, differences in evolutionary rates between temperate and tropical regions may explain the observed discrepancy in the dispersal-range size relationship. It has been argued that higher temperatures in the tropics accelerate evolutionary and ecological change (e.g., shorter generations times, fast mutation and selection rates), which may lead to increased speciation rates (Brown 2014). Higher speciation rates would negatively affect species range sizes, as new species rapidly arise, generally attaining a small range, and then give birth to new species again, thereby making them ‘ephemeral’ (Lester et al. 2007, Sheth et al. 2020). Therefore, in tropical regions there might be more species with small ranges independent of their dispersal abilities. This may be exacerbated by more narrow, specialized niches of tropical species that hinder the increase of range size and thus foster isolation, reproductive isolation and speciation (Janzen 1967). One way to circumvent this effect of evolution, is to consider species age (or evolutionary rates) directly when assessing the dispersal-range size relationship. Even though species age did not come out as a significant contributor in our analysis, we cannot discard its importance, because only 15 (1 in plants, 14 in animals) out of the 475 relationships considered species age.

Our results also suggested that studying multiple latitudes simultaneously prevents to find positive dispersal-range size relationships. This might be particularly relevant when combining latitudes for which the dispersal-range size relationship is different, like tropical vs. temperate regions together. Although a possible solution to circumvent this problem would be to correct for latitude, our results showed that correcting for latitude does not increase the chance of finding a positive dispersal-range size relationship. Nevertheless, only 30% of the relationships included a sort of latitude correction by either explicitly correcting for latitude (17 relationships) or restricting the study to a single latitudinal zone (107 relationships).

### The taxonomic effect on the dispersal-range size relationship

We found that studies that included species that can only be grouped into a high taxonomic level such as ‘phylum’ or ‘division’ found less often positive dispersal-range size relationships than studies that examined species within lower taxonomic levels (‘family’, ‘order’, ‘class’). Our separate analyses for plants and animals show that this effect is primarily driven by studies on plants, in which there was a clear distinction between e.g. ‘angiosperm-wide’ studies (126/159 relationships, in contrast to 10/298 relationships in animals that was at ‘phylum’ or higher taxonomic level), versus those performed for a specific ‘family’ or ‘genus’. This suggests that the disparity of dispersal-range size relationships when including plant (or animal) lineages that are phylogenetically distantly related and may therefore differ to a large extent in the traits that capture their dispersal ability (e.g., small seeds for wind-dispersed taxa, large seeds for animal-dispersed taxa, Table 2), will obscure or erase a relationship between dispersal and range size. In addition, including a clade in which many species are missing may also simply bias the initial dataset towards e.g. well-sampled species with relatively large range sizes. This means that a clear hypothesis and expectation on how dispersal affects range size in a studied clade is essential. In addition, including phylogenies to control for clade differences could provide a solution. We found that only 24% and 44% of dispersal-range size relationships that have been performed on animals and on plants, respectively, explicitly considered phylogenetic relationships. Even though phylogenetic correction was not selected in our final statistical model, we should be aware that we may not have had the statistical power to detect its real effect.

### Dispersal-range size moderators in animals vs. plants

Our results confirmed our hypothesis that a positive dispersal-range size relationship is less common for endotherms than for ectotherms (Table 1, Fig. 3, Table S2 Supple. Mat.), possibly because endotherms are less affected by niche limitation than ectotherms due to broad thermal tolerances, allowing them to attain and persist in larger ranges more easily. Indeed, the ‘thermal plasticity hypothesis’ proposes that high metabolic rates increase thermal tolerance, and the ‘energy constraint hypothesis’ that due to higher and sustained levels of energy requirements, organisms with high metabolic rates need to forage farther and with lower densities, resulting in larger home ranges and range sizes (Pie et al. 2021). Even though endo- and ectotherms might not differ in the dispersal capacities required during the transfer phase of dispersal, endothermy might have an advantage during settlement and establishment (the third phase of dispersal). If endothermy has allowed species to attain range sizes that are larger than predicted by their dispersal ability only, this might explain why we find fewer positive relationships between dispersal and range size. Note that a model including latitude instead of temperature regulation has a similarly good model fit (Table S1, S2 Suppl. Mat).

### Final remarks

In less than a century, macroecology has shed light on our understanding of species distribution patterns and about the processes and mechanisms governing them, at least from a theoretical point of view. Nowadays, we know that dispersal is a central process driving macroecological and macroevolutionary patterns in complex ways (e.g., Onstein et al. 2017, Alzate et al. 2019c, Sheard et al. 2020), but we still have not reached a consensus, based on empirical evidence, whether dispersal will positively affect geographical range sizes, or whether other variables, such as physiological tolerance (Pie et al. 2021), are more relevant for range size expansion, or even interact with aspects of the dispersal process (as shown here for endo- vs. ectotherms). Here, we show that differences between studies are largely responsible for different dispersal-range size relationship outcomes, which leads us to the following conclusions.

First of all, we need a better understanding of the dispersal process and envision it as a three-stage process (departure, transfer and settlement), with multiple traits (morphological, behavioral, physiological or life-history) acting differently in each of these stages (Clobert el al. 2012, Laube et al. 2013). It is important to be aware that many dispersal-related traits, which lead to net displacement, may be selected for functions other than dispersal *per se* (Burgess et al. 2005). In benthic marine organisms, dispersal-related traits (e.g., pelagic larvae, spawning mode) are often a by-product of traits selected for feeding, as part of the egg size-number trade-off, predation avoidance or retention of propagules (Burgess et al. 2015). In plants dispersal related traits, like seed size and plant height, can evolve as a result of the seed size-seed number trade-off and as to avoid competition for sunlight (Burgess et al. 2015). In animals, for which one type of locomotion is used for several ecological functions, dispersal can result from movement for foraging, exploration, mate and shelter seeking (Burgess et al. 2015). Regardless of whether traits are selected for dispersal or are an eco-evolutionary by-product, using a more complete picture of dispersal will allows us to capture its complexity in a more realistic manner and to better explain species geographical ranges. We should aim to use multiple dispersal-related traits, dispersal syndromes or co-variations of multiple dispersal-related traits (a multivariate dispersal phenotype) instead of using individual traits as dispersal proxies (Ronce & Clobert 2012).

Second, we propose ‘evolution’ and past dynamics as the fourth stage of the dispersal-range size paradigm. There is a lack of integration between macroecology and macroevolution (McGill et al. 2019); paleontological studies have pointed out the intricate relationship between dispersal, range size, speciation, extinction and species ages (Jablonski 1986), but very few studies have considered time for dispersal and range expansion (e.g., species age) or speciation/extinction dynamics when examining determinants of range size. Possibly, this is because suitably large and complete phylogenies, and reliable molecular clock models to estimate diversification rates, have been lacking until recently. In addition, past changes in paleoclimates, landscape connectivity, orogeny, and barriers, all have major impacts on dispersal and range size (Hagen et al. 2021), and should ideally be considered to fully capture this fourth temporal dimension to the dispersal-range size relationship. Finally, understanding the distribution of ranges, niche widths (e.g., right or left skewed), level of specialization, and distribution of niche properties (i.e., ecological opportunity for range expansion) within a studied system/clade, may allow the dissection of ecological and evolutionary processes influencing the dispersal-range size relationship.

## Acknowledgments

The authors acknowledge the support of iDiv funded by the German Research Foundation (DFG FZT 118, 202548816), specifically funding through sDiv, the Synthesis Centre of iDiv. We thank sDiv members, specially Roberto Rozzi, Alex Zizka and Juan Carvajal for lively discussion during this project. We thank Dries Bonte and Fons van der Plas for their useful comments and feedbacks on a previous version of this manuscript.

Map author: By Petr Dlouhý - World Map Blank.svg, Public Domain, https://commons.wikimedia.org/w/index.php?curid=984944

## References

1. Alzate, A., van der Plas, F., Zapata, F.A., Bonte, D. & Etienne, R.S. 2019a. Incomplete datasets obscure associations between traits affecting dispersal ability and geographic range size of reef fishes in the Tropical Eastern Pacific. Ecology and Evolution. 9: 1567– 1577.

2. Alzate, A., Etienne, R.S. & Bonte, D. 2019b. Experimental island biogeography demonstrates the importance of island size and dispersal for the adaptation to novel habitats. Global Ecology and Biogeography. 28: 238– 247.

3. Alzate, A, Janzen, T, Bonte, D, Rosindell, J, Etienne, RS. 2019c. A simple spatially explicit neutral model explains the range size distribution of reef fishes. Global Ecol Biogeogr. 28: 875– 890.

4. Blackburn, T.M. & Gaston, K.J. 1998. Some methodological issues in macroecology. The American Naturalist. 15(1): 68–83.

5. Böhning-Gaese, K., Caprano, T., van Ewijk, K. & Veith, M. 2006. Range Size: Disentangling Current Traits and Phylogenetic and Biogeographic Factors. Am. Nat. 167(4): 555–567.

6. Bonte, D., Van Dyck, H., Bullock, J.M., Coulon, A., Delgado, M., Gibbs, M., Lehouck, V., Matthysen, E., Mustin, K., Saastamoinen, M., Schtickzelle, N., Stevens, V.M., Vandewoestijne, S., Baguette, M., Barton, K., Benton, T.G., Chaput-Bardy, A., Clobert, J., Dytham, C., Hovestadt, T., Meier, C.M., Palmer, S.C.F., Turlure, C. & Travis, J.M.J. 2012. Costs of dispersal. Biological Reviews. 87: 290–312

7. Brown. J.H. 2014. Why are there so many species in the tropics? Journal of Biogeography. 41(1): 8–22.

8. Brown, J.H. & Kondric-Brown, A. 1977. Turnover rates in insular biogeography: effects of immigration on extinction. Ecology. 58: 445–449.

9. Brown, J.H., Stevens, G.C., & Kaufman, D.M. 1996. The geographic range: size, shape, boundaries, and internal structure. Annual Review of Ecology and Systematics. 27(1): 597–623

10. Burgess, S. C., Baskett, M. L., Grosberg, R. K., Morgan, S. G. & Strathmann, R. R. 2015. When is dispersal for dispersal? Unifying marine and terrestrial perspectives. Biological Reviews, 91(3), 867-882.

11. Chichorro, F., Juslén, A. & Cardoso, P. 2019. A review of the relation between species traits and extinction risk. Biological Conservation. 237:220–229.

12. Clobert, J., Baguette, M., Benton, T.G. & Bullock, J.M. 2012. Dispersal ecology and evolution. Oxford University Press. 462 p.

13. Dorie, V., Bates, D., Maechler, M., Bolker, B. & Walker, S. 2021. Bayesian Linear Mixed-Effects Models. V1.0-5. https://github.com/vdorie/blme

14. Fox, J. & Weisberg, S. 2019. An R Companion to Applied Regression, Third edition. Sage, Thousand Oaks CA. https://socialsciences.mcmaster.ca/jfox/Books/Companion/.

15. Gaston, K.J. & Blackburn, T.M. 1996. Range size - body size relationships: evidence of scale dependence. Oikos. 75(3):479–485.

16. Gaston, K. J. 2003. The structure and dynamics of geographic ranges. Oxford University Press, Oxford.

17. Gaston, K.J. & Fuller, R.A. 2009. The sizes of species’ geographic ranges. Journal of Applied Ecology, 46: 1–9.

18. Gaston, K.J. 2009. Geographic range limits: achieving synthesis. Proc. R. Soc. B. 276: 1395–1406.

19. Green, A.J., Baltzinger, C. and Lovas-Kiss, Á. (2021), Plant dispersal syndromes are unreliable, especially for predicting zoochory and long-distance dispersal. Oikos. https://doi.org/10.1111/oik.08327

20. Haffer, J. 1969. Speciation in Amazonian forest birds. Science. 165(3889): 131–137.

21. Hagen O, Flück B, Fopp F, Cabral JS, Hartig F, et al. (2021) gen3sis: A general engine for eco-evolutionary simulations of the processes that shape Earth’s biodiversity. PLOS Biology 19(7): e3001340.

22. Hanski, I., Kouki, J. & Halkka, A. 1993. Three explanations of the positive relationship between distribution and abundance of species. In: Species Diversity in Ecological Communities: Historical and Geographical Perspectives (eds Ricklefs, R.E. & Schluter, D.). University of Chicago Press, Chicago, pp. 108–116.

23. Hayes, J. P., Feldman, C. R. & Araújo, M.B. 2018. Mass-independent maximal metabolic rate predicts geographic range size of placental mammals. Functional Ecology. 32: 1194–1202.

24. Holt, R.D. 2003. On the evolutionary ecology of species’ ranges. Evol. Ecol. Res. 5:159– 78

25. Holt, R.D., Keitt, T.H., Lewis, M.A., Maurer, B.A. & Taper, M.L. 2005. Theoretical models of species’ borders: single species approaches. Oikos. 108:18–27

26. Holt, R. D. & Gomulkiewicz, R. 1997. How does immigration influence local adaptation? A re-examination of a familiar paradigm. The American Naturalist, 149, 563–572.

27. Hufbauer, R.A., Szűcs, M., Kasyon, E., Youngberg, C., Koontz, M.J., Richards, C., Tuff, T. & Melbourne, B. A. 2015. Three types of rescue can avert extinction in a changing environment. Proceedings of the National Academy of Sciences. 112 (33): 10557–10562.

28. Jablonski, D. 1986. Larval ecology and macroevolution in marine invertebrates. Bulletin of Marine Science. 39(2): 565–587.

29. Janzen, D.H. 1967. Why mountain passes are higher in the tropics. The American Naturalist. 101(919): 233–249.

30. Johnson, T.F., Isaac, N.J.B., Paviolo, A. & González-Suárez, M. 2021. Handling missing values in trait data. Global Ecology and Biogeography. 30: 51–62.

31. Laube, I., Korntheuer, H., Schwager, M., Trautmann, S., Rahbek, C. & B?hning-Gaese, K. 2013. Towards and more mechanistic understanding of traits and range sizes. Global Ecology and Biogeography. 22: 233–241.

32. Lester, S. E. & Ruttenberg, B. I. 2005. The relationship between pelagic larval duration and range size in tropical reef fishes: a synthetic analysis. Proceedings of the Royal Society B, 72, 585–591.

33. Lester, S. E., Ruttenberg, B. I., Gaines, S. D. & Kinlan, B. P. 2007. The relationship between dispersal ability and geographic range size. Ecology Letters, 10, 745–758.

34. Luiz, O. J., Allen, A. P., Robertson, D. R., Floeter, S. R., Kulbicki, M., Vigliola, L., Becheler, R. & Madin, J. S. 2013. Adult and larval traits as determinants of geographic range size among tropical reef fishes. Proceedings of the National Academy of Sciences, 110(41), 16498–16502.

35. Matthysen, E. 2012. Multicausality of dispersal: a review. In: Clobert, J., Baguette, M., Benton, T.G. & Bullock, J.M. 2012. Dispersal ecology and evolution. Oxford University Press. pp 3–18.

36. McGill, B. J., Chase, J. M., Hortal, J., Overcast, I., Rominger, A. J., Rosindell, J., et al. (2019). Unifying macroecology and macroevolution to answer fundamental questions about biodiversity. Glob. Ecol. Biogeogr. 28(12):1925–1936.

37. Mora, C., Treml, E., Robert. J., Crosby, K., Roy, D. & Titterson, D. P. 2012. High connectivity among habitats precludes the relationship between dispersal and range size in tropical reef fishes. Ecography, 35(1), 89–96.

38. Olson, D.M., E. Dinerstein, E.D. Wikramanayake, N.D. Burgess, G.V.N. Powell, E.C. Underwood, J.A. D’Amico, I. Itoua, H.E. Strand, J.C. Morrison, C.J. Loucks, T.F. Allnutt, T.H. Ricketts, Y. Kura, J.F. Lamoreux, W.W. Wettengel, P. Hedao, & K.R. Kassem. 2004. Terrestrial Ecoregions of the World: A New Map of Life on Earth (PDF, 1.1M) BioScience 51:933–938.

39. Onstein, R.E., Baker, W.J., Couvreur, T.L.P. et al. 2017. Frugivory-related traits promote speciation of tropical palms. Nat Ecol Evol 1, 1903–1911.

40. Onstein, R.E., Kissling, W.D., Chatrou, L.W., Couvreur, T.L.P., Morlon, H. & Sauquet, H. 2019. Which frugivory-related traits facilitated historical long-distance dispersal in the custard apple family (Annonaceae)? Journal of Biogeography. 46: 1874– 1888.

41. Pie, M. R., Divieso, R. & Caron, F. S. 2021. Do geographic range sizes evolve faster in endotherms? Evolutionary Biology. https://doi.org/10.1007/s11692-021-09537-x

42. Purvis, A., Gittleman, J. L., Cowlishaw, G. & Mace, G. M. 2000. Predicting extinction risk in declining species. Proc. R. Soc. Lond. B. 267(1456):1947–1952.

43. Rangel, T.F.L.V.B. & Diniz-Filho, J.A.F. 2005. Neutral community dynamics, the mid-domain effect and spatial patterns in species richness. Ecology Letters. 8: 783–790.

44. Ronce, O. 2007. How does it feel to be like a rolling stone? Ten questions about dispersal evolution. Ann. Rev. Ecol. Evol. Syst. 38:231–253.

45. Ronce, O. & Clobert, J. 2012. Dispersal syndromes. In: Clobert, J., Baguette, M., Benton, T.G. & Bullock, J.M. (eds). Dispersal ecology and evolution. Oxford University Press. pp 119–138.

46. Schurr, F.M., Midgley, G.F., Rebelo, A.G., Reeves, G., Poschlod, P. & Higgins, S.I. 2007. Colonization and persistence ability explain the extent to which plant species fill their potential range. Global Ecology and Biogeography. 16: 449–459.

47. Sexton, J. P., McIntyre, P. J., Angert, A. L. & Rice, K. J. 2009. Evolution and ecology of species range limits. Annual Review of Ecology, Evolution and Systematics, 40, 415–436.

48. Sheard, C., Neate-Clegg, M.H.C., Alioravainen, N., Jones, S.E.I., Vincent, C., MacGregor, H.E.A., Bregman, T.P., Claramunt, S. & Tobias, J. A. 2020. Ecological drivers of global gradients in avian dispersal inferred from wing morphology. Nature Communications. 11: 2463.

49. Sheth, S.N., Morueta-Holme, N., Angert, A.L. 2020. Determinants of geographic range size in plants. New Phytologist. 226:650–665.

50. Spalding, M.D., Fox, H.E., Allen, G.R., Davidson, N., Ferdaña, S.A., Finlayson, M., Halpern, B.S., Jorge, M.A., Lombana, A., Lourie, S.A., Martin, K.D., McManus, E., Molnar, J., Recchia., C.A., & Robertson, J. 2007. Marine ecoregions of the world: a bioregionalization of coastal and shelf areas. BioScience. 57(7): 573–583.

51. Storch, D. & Šizling, A.L. 2008. The concept of taxon invariance in ecology: do diversity patterns vary with changes in taxonomic resolution? Folia Geobotanica. 43(3): 329–344.

52. Svenning, J.-C. & Skov, F. 2004. Limited filling of the potential range in European tree species. Ecology Letters, 7: 565–573.

53. Svenning, J.-C., Normand, S. & Skov, F. 2008. Postglacial dispersal limitation of widespread forest plant species in nemoral Europe. Ecography. 31: 316–326.

54. Thieltges, D.W., Hof, C., Borregaard, M.K., Dehling, M., Brändle, M., Brandl, R. & Poulin, R. 2011. Range size patterns in European freshwater trematodes. Ecography. 34: 982–989.

55. Travis, J.M.J., Mustin, K., Bartoń, K.A., Benton, T.G., Clobert, J., Delgado, M.M., Dytham, C., Hovestadt, T., Palmer, S.C.F., Van Dyck, H. and Bonte, D. (2012), Modelling dispersal: an eco-evolutionary framework incorporating emigration, movement, settlement behaviour and the multiple costs involved. Methods in Ecology and Evolution. 3: 628–641.

56. Varadhan, R., Borchers, H.W. & Bechard, V. 2020. dfoptim: Derivative-Free Optimization. R package version 2020.10-1.

57. Webb, T. J., & Gaston, K. J. (2000). Geographic range size and evolutionary age in birds. Proc. R. Soc. B. 267(1455): 1843–1850.

58. Willis, J.C. 1922. Age and Area: a study in geographical distribution and origin of species. Cambridge: Cambridge University Press.

59. Zapata, F. A. & Herrón, P. A. (2002) Pelagic larval duration and geographic distribution of tropical eastern Pacific snappers (Pisces: Lutjanidae). Marine Ecology Progress Series, 230, 295–300.

